# Morphological profile determines the frequency of spontaneous calcium events in astrocytic processes

**DOI:** 10.1101/410076

**Authors:** Yu-Wei Wu, Susan Gordleeva, Xiaofang Tang, Pei-Yu Shih, Yulia Dembitskaya, Alexey Semyanov

## Abstract

Astrocytes express a complex repertoire of intracellular Ca^2+^ transients (events) that represent a major form of signaling within individual cells and in the astrocytic syncytium. These events have different spatiotemporal profiles, which are modulated by neuronal activity. Spontaneous Ca^2+^ events appear more frequently in distal astrocytic processes and independently from each other. However, little is known about the mechanisms underlying such subcellular distribution of the Ca^2+^ events. Here we identify the initiation points of the Ca^2+^ events within the territory of single astrocytes expressing genetically encoded Ca^2+^ indicator GCaMP2 in culture or in hippocampal slices. We found that most of the Ca^2+^ events start in thin distal processes. Our mathematical model demonstrated that a high surface-to-volume (SVR) of the thin processes leads to increased amplitude of baseline Ca^2+^ fluctuations caused by a stochastic opening of Ca^2+^ channels in the plasma membrane. Suprathreshold fluctuations trigger Ca^2+^-induced Ca^2+^ release (CICR) from the Ca^2+^ stores by activating inositol 1,4,5-trisphosphate (IP_3_) receptors. In agreement with the model prediction, the spontaneous Ca^2+^ events frequency depended on the extracellular Ca^2+^ concentration. Astrocytic depolarization by high extracellular K^+^ increased the frequency of the Ca^2+^ events through activation of voltage-gated Ca^2+^ channels (VGCC) in cultured astrocytes. Our results suggest that the morphological profile of the astrocytic processes is responsible for tuning of the Ca^2+^ event frequency. Therefore, the structural plasticity of astrocytic processes can be directly translated into changes in astrocytic Ca^2+^ signaling. This may be important for both physiological and pathological astrocyte remodeling.

**Main points:**

- Majority of spontaneous Ca^2+^ events start in thin astrocytic processes
- Higher surface-to-volume ratio of the process is responsible for larger intracellular Ca^2+^ fluctuations
- Larger intracellular Ca^2+^ fluctuations trigger Ca^2+^-dependent Ca^2+^ release

## 1. Introduction

Ca^2+^ activity is a major form of signaling in astrocytes (Agulhon et al., 2008; Bazargani & Attwell, 2016; Rusakov, Zheng, & Henneberger, 2011; Verkhratsky, Orkand, & Kettenmann, 1998). Changes in intracellular Ca^2+^ ([Ca^2+^]_i_) are required for many astrocytic functions, including release of gliotransmitters (Araque et al., 2014; Zorec et al., 2012), regulation of local blood flow (Petzold & Murthy, 2011), regulation of K^+^ uptake (F. Wang et al., 2012), morphological plasticity of astrocytic processes (Heller & Rusakov, 2015; Tanaka et al., 2013). Large and slow astrocytic Ca^2+^ events are thought to be predominantly mediated by the metabotropic pathways linked to inositol 1,4,5-trisphosphate (IP_3_) production, which triggers CICR through IP_3_ receptors (IP_3_Rs) (De Young & Keizer, 1992; Kirischuk, Kirchhoff, Matyash, Kettenmann, & Verkhratsky, 1999; Ullah, Jung, & Cornell-Bell, 2006). Stimulation of Shaffer collaterals or an application of metabotropic glutamate receptor (mGluR) agonists increase the frequency and the spreading of the astrocytic Ca^2+^ events in hippocampal slices (Haustein et al., 2014; Sun et al., 2014; Wu et al., 2014). The blockers of mGluRs or tetrodotoxin (TTX, blocks neuronal action potentials) reduce the frequency and the spread of the spontaneous Ca^2+^ events in astrocytes (Haustein et al., 2014; Rungta et al., 2016; Sun et al., 2014; Wu et al., 2014). However, they fail to block astrocytic Ca^2+^ activity completely (Haustein et al., 2014; Rungta et al., 2016; Sun et al., 2014; Wu et al., 2014). Indeed, Ca^2+^ can enter astrocytes through ionotropic receptors in plasma membrane (Fumagalli et al., 2003; Palygin, Lalo, Verkhratsky, & Pankratov, 2010; C. M. Wang, Chang, Kuo, & Sun, 2002). Yet, the spontaneous Ca^2+^ events are still detectable in astrocytes in the presence of AMPA, NMDA, GABA_B_, P_2_X, and P_2_Y receptor blockers (Rungta et al., 2016; Wu et al., 2014). Thus, at least a fraction of the spontaneous Ca^2+^ events in astrocytes is not triggered by activation of these receptors.

In fact, astrocytes possess several other mechanisms for the Ca^2+^ entry through the plasma membrane. Expression of VGCCs has been reported in astrocytes both in culture (Latour, Hamid, Beedle, Zamponi, & Macvicar, 2003; Parpura, Grubisic, & Verkhratsky, 2011; Yaguchi & Nishizaki, 2010) and in hippocampal slices (Letellier et al., 2016). Activation of L-type VGCCs upon astrocyte depolarization is involved in the release of gliotransmitters and regulation of presynaptic strengths (Letellier et al., 2016; Yaguchi & Nishizaki, 2010). However, the Ca^2+^ entry through the VGCCs may still need to be amplified by CICR (Carmignoto, Pasti, & Pozzan, 1998). Na^+^/Ca^2+^ exchanger (NCX) can transport Ca^2+^ into the astrocytes upon increase in intracellular Na^+^ concentration which occurs during glutamate uptake (Brazhe, Verisokin, Verveyko, & Postnov, 2018; Oschmann, Mergenthaler, Jungnickel, & Obermayer, 2017; Rojas et al., 2007). Other sources of the Ca^2+^ entry in astrocytes include store-operated channels (SOCs) ORAI1 and transient receptor potential type A channels (TRPAs) (Bazargani & Attwell, 2016; Shigetomi, Jackson-Weaver, Huckstepp, O’Dell, & Khakh, 2013).

Notably, the majority of Ca^2+^ events start in fine distal astrocytic processes (Asada et al., 2015; Bindocci et al., 2017; Nakayama, Sasaki, Tanaka, & Ikegaya, 2016; Nett, Oloff, & McCarthy, 2002). This can be attributed to the location of synapses (Di Castro et al., 2011) and subcellular distribution of receptors, channels and transporters in astrocytes (Arizono et al., 2012; Hayashi & Yasui, 2015). Here we show that spontaneous Ca^2+^ events can start in thin astrocytic processes because of their high SVR: Ca^2+^ entry though plasma membrane produces larger elevations in [Ca^2+^]_I_ that more likely to trigger CICR.

## 2. Materials and Methods

### 2.1 Primary Hippocampal Astrocyte/Neuron Co-culture and Transfection

Primary hippocampal astrocyte/neuron co-cultures were prepared from Wistar rats (Japan SLC Inc.) at embryonic day 18–20 with slight modifications of the previously described procedure (Wu et al., 2014). All procedures were performed in accordance with RIKEN regulations. Briefly, hippocampi from 6 to 8 pups were dissociated in petri dishes filled with Hank’s balanced salt solution (HBSS) containing 20 mM N-2-hydroxyethyl-piperazine-N’-2-ethane-sulfonic acid (HEPES). The dissected hippocampi were digested by incubation in 3 ml HBSS containing 20 mM HEPES, 0.125% trypsin, and 0.025% DNase I for 5 min at 37°C. After incubation, the pieces were washed 3 times with HBSS containing 20 mM HEPES and triturated with pipettes in the plating medium containing Minimal Essential Medium, B27, glutamine, sodium pyruvate, and penicillin-streptomycin. The cells were plated at a density of 1.2–1.4 x 10^5^ cells/well in culture plates containing 18-mm glass coverslips coated with 0.04% polyethyleneimine (Sigma-Aldrich, St. Louis, MO). Primary cell cultures were maintained at 37°C in a 5% CO_2_ humid incubator until used for experiments at 7 to 13 days *in vitro*. Every 3d day, half of the volume of the medium was replaced with Neurobasal medium containing B27, glutamine, and penicillin-streptomycin. These procedures lead astrocytes to develop the complex shapes without a need for serum-free base medium as previously suggested (Liddelow et al., 2017). For transfection, a coverslip in 1 ml culture medium was supplied with the transfection mixture containing 100 µl OPTI-MEM (Invitrogen), 0.5 µg GCaMP2 DNA plasmid, and 1 µl Lipofectamine 2000 Transfection Reagent (Invitrogen). The culture was incubated with this mixture for 24 hours before imaging.

### 2.2 Ca^2+^ Imaging in Cultured Astrocytes

Ca^2+^ imaging of GCaMP2-expressing cultured astrocytes was performed in a recording chamber where the cultures were continuously superfused with a balanced salt solution containing (in mM): 115 NaCl, 5.4 KCl, 2 CaCl_2_, 1 MgCl_2_, 10 D-glucose, and 20 HEPES (pH 7.4, 33-34°C). Fluorescence was detected using an inverted microscope (IX70, Olympus) equipped with an objective lens Plan Apo 60x, NA 1.42 (Olympus) and a cooled charge-coupled device camera (array size: 256 x 337 pixels, 0.174 µm^2^/pixel; ORCA II-ER, Hamamatsu Photonics). A light source, a 490-nm light-emitting diode illumination system (precisExcite, CoolLED), and appropriate filter sets (470–490 nm for excitation; 515–550 nm for emission) were used. Images were acquired at 2 Hz using MetaView software (Meta Imaging, Dowington, PA). Spontaneous Ca^2+^ activity was recorded for 20 min in the presence of a cocktail of blockers for voltage-gated Na^+^ channels (TTX, 1µM), mGluRs ((S)-α-Methyl-4-carboxyphenylglycine (S-MCPG), 100 µM), NMDA receptors (D-2-amino-5-phosphonovalerate (D-AP5), 25 µM), AMPA receptors (2,3-Dioxo-6-nitro-1,2,3,4-tetrahydrobenzo[f]quinoxaline-7-sulfonamide (NBQX), 25 µM), GABA_B_ receptors (3-[[(3,4-Dichlorophenyl)methyl]amino]propyl] diethoxymethyl)phosphinic acid (CGP 52432), 5 µM), and purinergic P2 receptors (Pyridoxalphosphate-6-azophenyl-2’,4’-disulfonic acid (PPADS), 30 µM). In experiments with different extracellular Ca^2+^ concentrations (2, 3, and 4 mM) imaging was performed during 10 min for each concentration.

### 2.3 Estimation of Surface-to-Volume Ratio

Morphology of the cultured astrocytes was reconstructed from baseline fluorescence images of GCaMP2. Imaging was done with a laser scanning microscope FV-1000 (Olympus, Japan) equipped with a 488 nm laser and a 60x water immersion objective (NA 1.0). The heights of the astrocytic processes were estimated by the full-width-at-half-maximum (half-width) of the fluorescence signal along the z-axis (Figure 3A and inset). The conversion factor between the heights of the processes imaged with the confocal microscopy and fluorescence intensity of the same processes imaged with CCD camera was obtained from linear regression (*n* = 23 processes/3 cells; Figure 3D). The SVRs of the processes were then calculated as a triangle from the half-width (HW) and the height of the process as

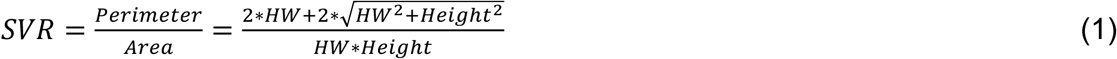

### 2.4 Mitochondria staining

The cultured astrocytes were treated with 50 nM MitoTracker Red CMXRos (Thermo Fisher Scientific Inc., USA) for 15 min at 37°C and then washed twice by culture medium and one time by Neurobasal medium (Thermo Fisher Scientific Inc., USA). The cultured cells were then fixed with 3.7% formaldehyde at room temperature for 10 min, followed by washing with PBS before mounting and imaging.

### 2.5 Two-photon Ca^2+^ imaging in hippocampal slices

The transverse hippocampal slices were prepared from 5 – 6 months-old double transgenic mice carrying tetO-GCaMP2 and GLT-1-tTA, which led to astrocyte-specific expression of GCaMP2 (Tanaka et al., 2013). The animals were kept and killed according to RIKEN regulations. The brains were removed, and the two hippocampi were dissected. The 350-µm thick slices were prepared with vibrating microtome Microm HM 650 V (Thermo Fisher Scientific Inc., USA) in an ice-cold cutting solution containing (in mM): 75 sucrose, 87 NaCl, 2.5 KCl, 0.5 CaCl_2_, 1.25 NaH_2_PO_4_, 7 MgCl_2_, 25 NaHCO_3_, 1 Na-ascorbate, and 11 D-glucose. Then the slices were transferred to an interfaced chamber with a storage solution containing (in mM): 127 NaCl, 2.5 KCl, 1.25 NaH_2_PO_4_, 1 MgCl_2_, 1 CaCl_2_, 25 NaHCO_3_, and 25 D-glucose, and kept at room temperature for at least 1 h prior imaging. Then the slices were transferred to an imaging chamber and superfused with a Ringer solution containing (in mM): 127 NaCl, KCl, 1.25 NaH_2_PO_4_, 1 MgCl_2_, 2 CaCl_2_, 25 NaHCO_3_, and 25 D-glucose at 34°C. All solutions were saturated with 95% O_2_ and 5% CO_2_ gas mixture. The osmolarity was adjusted to 300 mOsm. The astrocyte-specific expression of GCaMP2 was sparse, therefore, individual astrocytes could be imaged. Two-photon time-lapse imaging (1 frame per second) was done with the laser scanning microscope FV-1000 (Olympus, Japan). GCaMP2 was excited at 890 nm wavelength with a femtosecond laser Chameleon XR (Coherent, USA) and the fluorescence was collected with a 60x water immersion objective (NA 1.0) and a 495–540 nm bandpass filter.

### 2.6 Drugs and Chemicals

All drugs were made from 1000x stock solutions that were kept frozen at −20°C in 100-200 µl aliquots. CGP 52432, TTX, S-MCPG, PPADS, NBQX, HC 030031, and D-AP5 were purchased from Tocris Cookson (Bristol, UK). The astrocytic marker, Sulforhodamin 101, was purchased from Thermo Fisher Scientific Inc. (USA), and all other drugs and chemicals were purchased from Sigma-Aldrich (USA).

### 2.7 Data Analysis and Statistics

Whole Ca^2+^ events were automatically identified with custom-made software (Matlab, Mathworks) and analyzed following the algorithm described by Wu et al., 2014. In brief, the events were defined by combining two different thresholds: a dynamic threshold and an absolute threshold. The dynamic threshold was set at two standard deviations of baseline fluorescence fluctuations in each pixel. The absolute threshold was ΔF/F = 0.1. If the neighboring ‘active’ pixels passed the absolute threshold and contained areas detected by the dynamic threshold, the areas were defined as the Ca^2+^ event area in a particular frame. The event areas were then connected in the subsequent frames forming thus whole Ca^2+^ event reconstructed in x-y-time 3D space (Figure 2C). Two Matlab (version 2015a) functions were used to find the weighted centroid of the first frame of each whole Ca^2+^ event, defined as the initiation point: “*bwconncomp*” and “*regionprops*”. The function “*regionprops*” returned the coordinates specifying the center of the region based on location and intensity value (Figure 2D).

**Figure 1.**
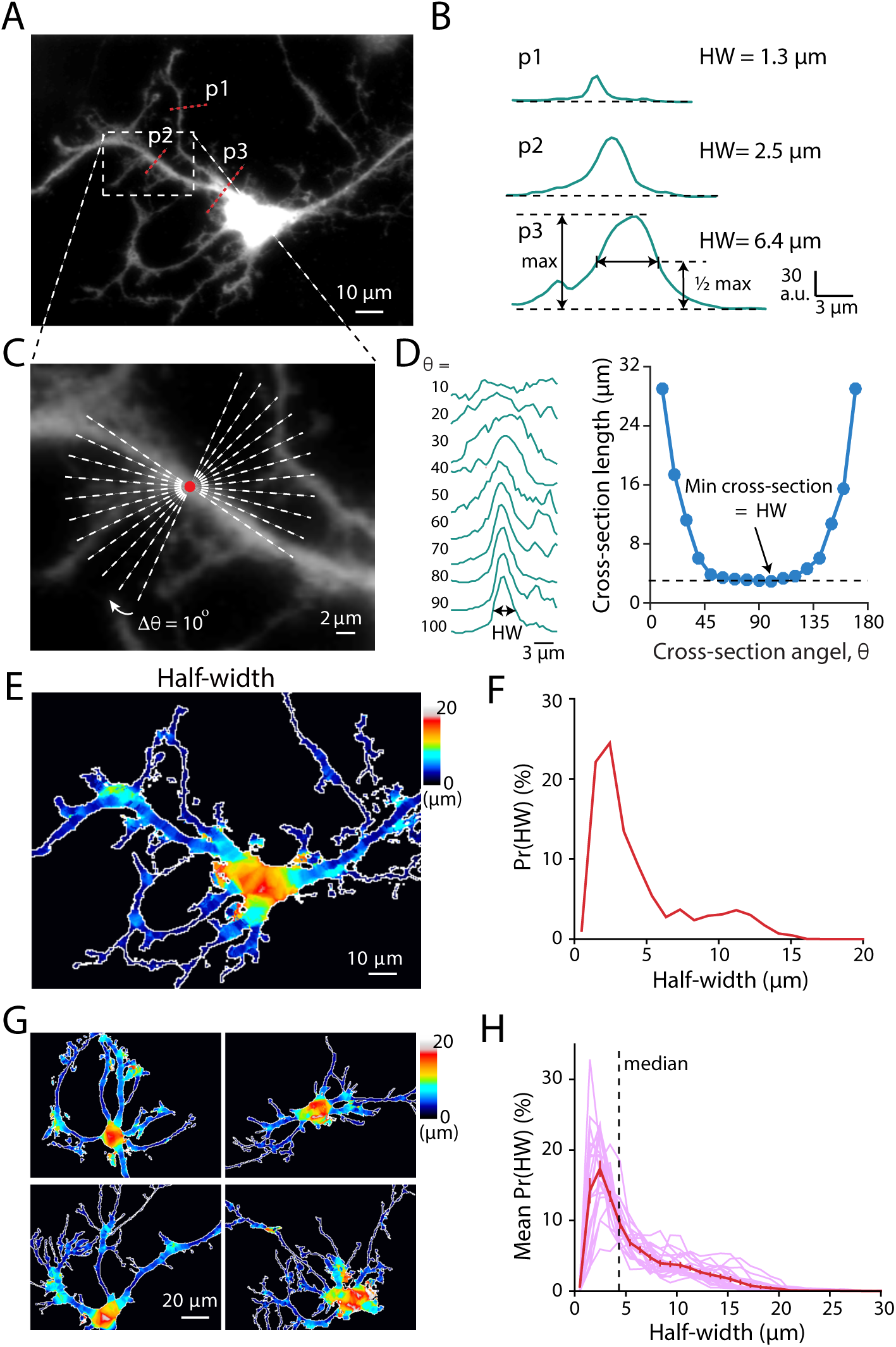
Morphological profile of cultured hippocampal astrocyte. **(A)** A typical astrocyte in culture expressing the Ca^2+^ sensor GCaMP2. **(B)** Fluorescent profiles of three astrocytic processes of different sizes (p1, p2, and p3) marked in (A). The sizes of the processes were defined by their half-width (HW). **(C and D)** Determining the local HW of astrocytic processes with the minimum cross-section. An enlarged image of astrocytic processes from the boxed area in (A). For each pixel within the astrocytic territory, 18 cross-sections were made with a 10° increment (white dash-lines). **(D)** *Left*, representative examples of cross-sections defined in (C). *Right*, the HW of the process for each pixel was defined as the minimum cross-section length. **(E)** The local HW of the astrocyte is indicated with the color map. Colder color indicates thinner processes. **(F)** The probability density distribution of HWs (Pr(HW)) in the astrocyte in (E). **(G)** Four examples of the HW color maps. **(H)** The mean Pr(HW) of several astrocytes (thin lines – individual cells, n = 22 cells, thick line mean ± SEM). Dashed line indicates the median (4.5 ± 0.3 µm) of the HW distribution.

**Figure 2.**
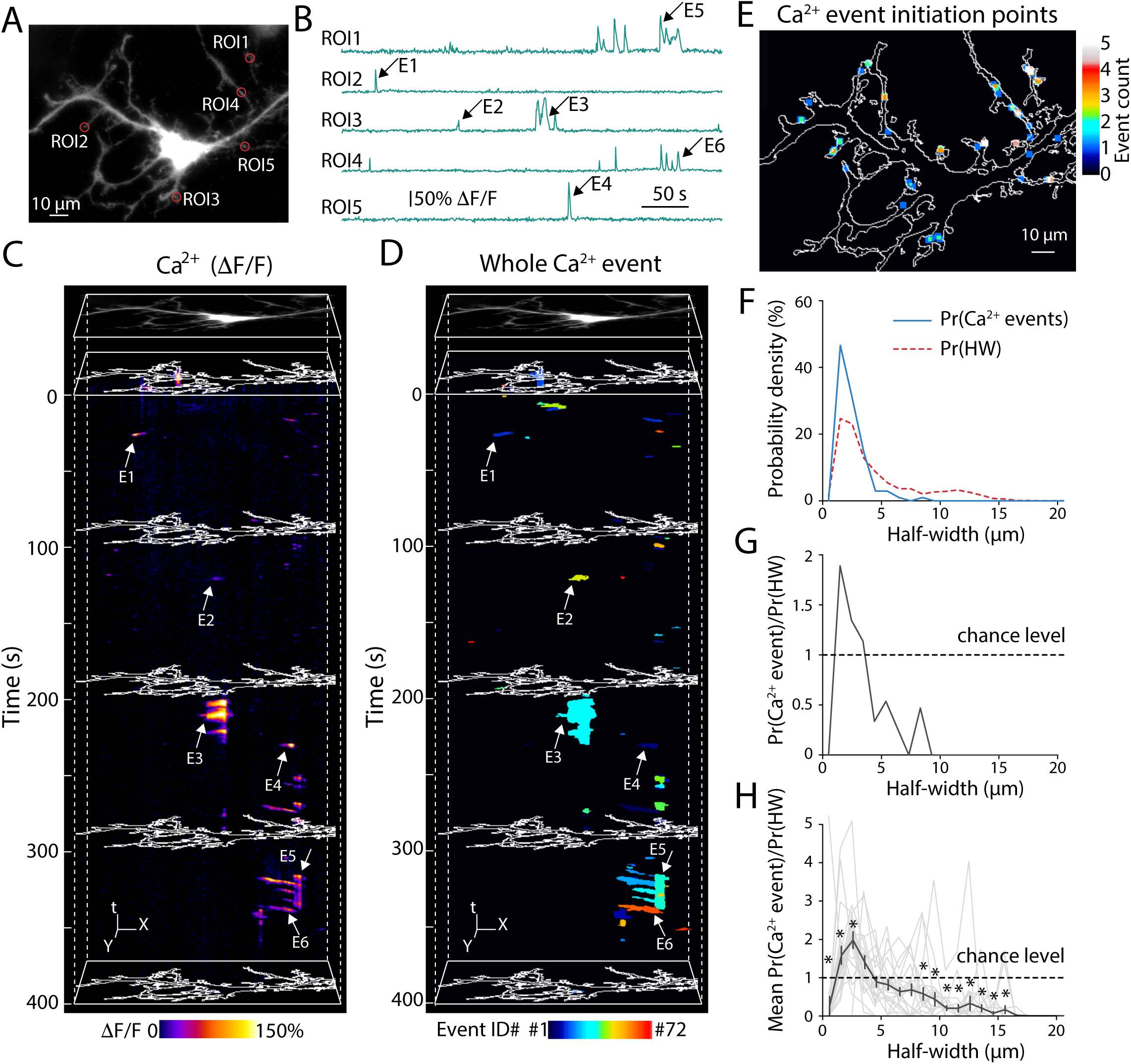
Ca^2+^ events predominantly start in thinner processes of astrocytes. **(A)** An image of cultured astrocyte expressing GCaMP2. **(B)** Ca^2+^ dynamics in the region-of-interests (ROIs) indicated in (A). **(C)** A representative example of X-Y-time heat map of Ca^2+^ dynamics (ΔF/F). Arrows indicate six examples of Ca^2+^ transients (E1-E6) shown in ROI-based traces in (B). The white contour shows the boundaries of the astrocyte. **(D)** A representative example of X-Y-time binary map of detected whole Ca^2+^ events. Random colors were assigned to individual events to distinguish their boundaries. The same examples of the detected whole Ca^2+^ events were indicated with arrows. **(E)** A representative image of initiation points of Ca^2+^ events. The centroids of the initial frame of 72 Ca^2+^ events were plotted, and the counts were presented as a color map. **(F)** A representative example of the probability density of Ca^2+^ events initiation at particular HWs (Pr(Ca^2+^ events), blue line) from the astrocyte in (E). Dashed pink line indicates Pr(HW) as in Figure 1F. **(G)** The ratio Pr(Ca^2+^ events)/Pr(HW) is equal to 1 when the probability of events initiation at particular HW is determined by the probability of such HW in the astrocyte (chance level). The ratio >1 suggests that Ca^2+^ events appear in the processes with this HW above chance level and vice versa for the ration <1. **(H)** The summary of the Pr(Ca^2+^ events)/Pr(HW) ratio (thin lines – individual cells, n = 22 cells, thick line - mean ± SEM). The Pr(Ca^2+^ events)/Pr(HW) ratio was significantly higher that the chance level in the thinner processes and significantly lower in the thicker processes and the soma. *p < 0.05 ANOVA post-hoc Bonferroni’s multiple comparisons.

Statistical significance was tested using Wilcoxon signed-rank test for paired comparison; Friedman test for equivalent repeated measurement (RM) one-way analysis of variance (ANOVA); two-way RM ANOVA followed by *post-hoc* Bonferroni’s multiple comparisons for the paired measurements across treatments and morphological parameters using Prism 7.0 (GraphPad, USA). The level of significance was set at *p* < 0.05. All group measures presented in the figures and in the text are given as mean ± the standard error of mean (SEM).

### 2.8 Modeling of Ca^2+^ Dynamics

Astrocytic [Ca^2+^]_i_ changed due to several types of effluxes and influxes considered in the model. Ca^2+^ was released to the cytosol from endoplasmic reticulum (ER) through IP_3_Rs (*J*_*IP3*_). Sarco/endoplasmic reticulum Ca^2+^-ATPase (SERCA) pumped Ca^2+^ back into the ER (*J*_*pump*_). In addition, we introduced a leak current from the ER to the cytosol (*J*_*leak*_). The influx and the efflux of Ca^2+^ through the plasma membrane were also considered (*J*_*in*_ and *J*_*out*_, respectively)

The balance of the Ca^2+^ fluxes to and from the cytosol was described by:

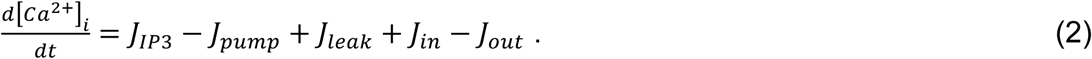

The kinetics of the IP_3_R mediated flux was based on the Li-Rinzel simplification (Y. X. Li & Rinzel, 1994) of the De Young-Keizer model (De Young & Keizer, 1992):

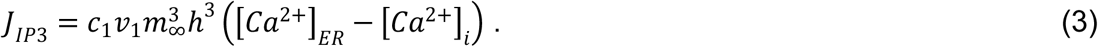

These kinetics were written analogously to the Hodgkin–Huxley model of the action potential (Hodgkin & Huxley, 1952). *c*_*1*_ is the ratio of the ER vs the cytosol volumes (Ullah et al., 2006). *v*_*1*_ is the maximal rate of the IP_3_-dependent CICR. The driving force for the Ca^2+^ fluxes was created by the concentration gradient between the ER ([Ca^2+^]_*ER*_) and the cytosol ([Ca^2+^]_i_).

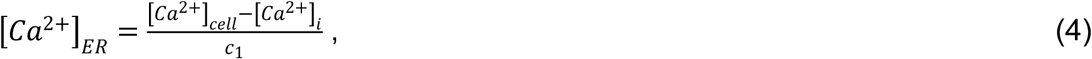

where [Ca^2+^]_*cell*_ is the total cellular Ca^2+^ concentration.

The activation Variable of the IP_3_R *m*_*∞*_ was described by

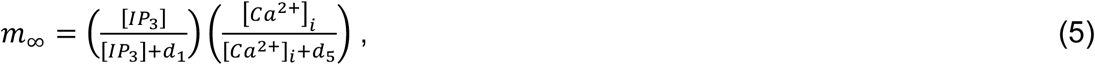

where [IP_3_] is the IP_3_ concentration in the cytosol, *d*_*1*_ is the dissociation constant for IP_3_, *d*_*5*_ is the Ca^2+^ activation constant (see Table 1).

**Table 1.** The parameter Values used in the model. The values of parameters for the Ca^2+^ fluxes through the membrane of astrocytes or through the membrane of the ER were taken from the previous studies (De Young & Keizer, 1992; B. Li, Chen, Zeng, Luo, & Li, 2012; Y. X. Li & Rinzel, 1994; Ullah et al., 2006; Zeng et al., 2009).

| Parameter | Definition | Value |
| --- | --- | --- |
| $[Ca^{2+}]_{cell}$ | The total cell free $Ca^{2+}$ concentration in the cell | 2.0 $\mu M$ |
| $c_1$ | (ER volume)/(cytosolic volume) | 0.185 |
| $v_1$ | Maximal CICR rate | 6.0 $s^{-1}$ |
| $v_2$ | Maximal rate of $Ca^{2+}$ leak from the ER | 0.11 $s^{-1}$ |
| $v_3$ | Maximal rate of SERCA uptake | 2.2 $\mu M s^{-1}$ |
| $J_{pass}$ | Rate of $Ca^{2+}$ leak across the plasma membrane | 0.025 $\mu M s^{-1}$ |
| $v_6$ | Maximal rate of activation-dependent $Ca^{2+}$ influx | 0.2 $\mu M s^{-1}$ |
| $k_1$ | Rate constant of $Ca^{2+}$ extrusion | 0.5 $s^{-1}$ |
| $k_2$ | Half-saturation constant for $Ca^{2+}$ entry | 1.0 $\mu M$ |
| $k_3$ | SERCA $Ca^{2+}$ affinity | 0.1 $\mu M$ |
| $a_2$ | IP<sub>3</sub>R binding rate for $Ca^{2+}$ inhibition | 0.14 $\mu M^{-1} s^{-1}$ |
| $d_1$ | Dissociation constant for IP<sub>3</sub> | 0.13 $\mu M$ |
| $d_2$ | $Ca^{2+}$ inactivation dissociation constant | 1.049 $\mu M$ |
| $d_3$ | Receptor dissociation constant for IP<sub>3</sub> | 0.9434 $\mu M$ |
| $d_5$ | $Ca^{2+}$ activation dissociation constant | 0.082 $\mu M$ |
| $\alpha$ | | 0.8 |
| $v_4$ | Maximal rate of IP<sub>3</sub> production by PLCδ | 0.4 $\mu M s^{-1}$ |
| $1/\tau_r$ | Rate constant for IP<sub>3</sub> degradation | 0.14 $s^{-1}$ |
| $[IP_3^*]$ | Steady state concentration of IP<sub>3</sub> | 0.25 $\mu M$ |
| $k_4$ | Inhibition constant of PLCδ activity | 1.1 $\mu M$ |
| Parameters for VGCCs |  |  |
| $g$ | Conductance density of VGCC | 3.5 pS $\mu m^{-2}$ |
| $r$ | Radius of an astrocytic process | 0.01-7 $\mu m$ |
| $l$ | Unitary length | 1 $\mu m$ |
| $\pi$ | | 3.14 |
| $R$ | Ideal gas constant | 8.31 J mol<sup>-1</sup> K<sup>-1</sup> |
| $T$ | Absolute temperature | 293 K |
| $F$ | Faraday constant | 96485 C mol<sup>-1</sup> |
| $z_K$ | Valence of K<sup>+</sup> | 1 |
| $z_{Ca}$ | Valence of Ca<sup>2+</sup> | 2 |
| $\epsilon$ | Modulation factor | 17 mV |
| $[Ca^{2+}]_o$ | Extracellular $Ca^{2+}$ concentration | 10<sup>-5</sup>- 4 mM |
| $\sigma_m^2$ | Variance of noise | 0.17 |

The inactivation variable of the IP_3_R *h* was modeled as a dynamic variable (Y. X. Li & Rinzel, 1994):

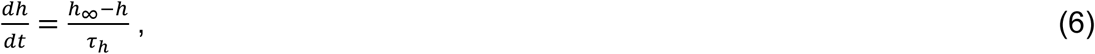

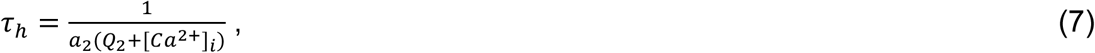

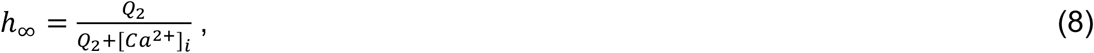

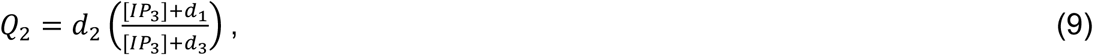

where *d*_*1*_ is the dissociation constant for IP_3_, *d*_3_ is the receptor dissociation constant for IP_3_ (see Table 1).

The SERCA mediated flux from the cytosol to the ER was described by the Hill-type kinetic model (Y. X. Li & Rinzel, 1994):

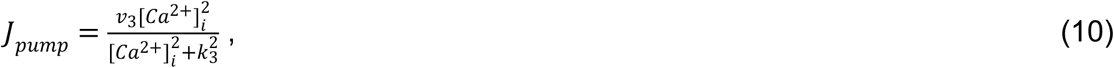

where *v*_3_ is maximal Ca^2+^ uptake and *k*_3_ is activation constant for ATP-Ca^2+^ pump.

The leak flux from the ER to the cytosol was described by:

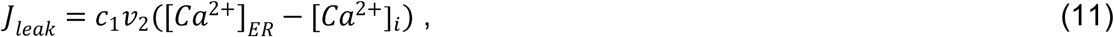

where *v*_*2*_ is the maximal leak flux through Ca^2+^ channels.

The Ca^2+^ influx through plasma membrane (*J*_*in*_) was described by the sum of 3 fluxes: the constant Ca^2+^ influx (*J*_*pass*_) through passive channels, the capacitive Ca^2+^ entry (*J*_*CCE*_), and the influx through VGCCs (*J*_*VGCC*_).

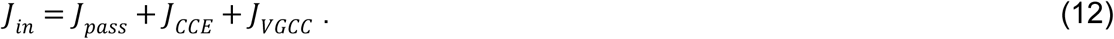

*J*_*CCE*_ depended on the IP_3_ concentration (Dupont & Goldbeter, 1993):

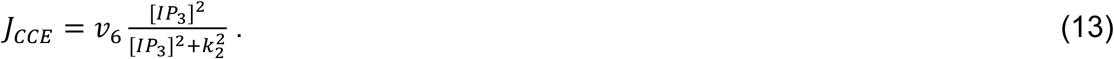

*J*_*VGCC*_ was calculated as described in (Zeng, Li, Zeng, & Chen, 2009):

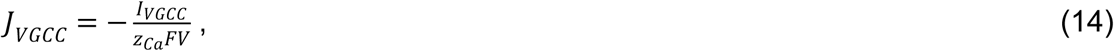

where *z*_*Ca*_ is the valence of Ca^2+^; *F* is the Faraday constant; *V* is the volume of an astrocytic compartment. Since we considered a chunk of an astrocytic process as a cylinder with radius *r* and unitary length *l = 1*, the volume of the process was: *V* = *πr*^*2*^*l = πr*_*2*_.

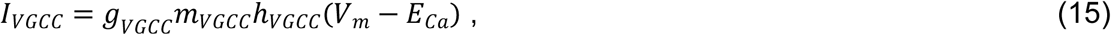

where *g*_*VGCC*_ is VGCC-mediated conductance. *g*_*VGCC*_ = *gs,* where *g* is the conductance density and *S* is the surface of the plasma membrane, which was equal to the side surface of the cylinder with unitary length *l* = 1: *S* = *2πrl = 2πr; m*_*VGCC*_ and *h*_*VGCC*_ are gates that regulate activation and inactivation of the VGCCs (Zeng et al., 2009):

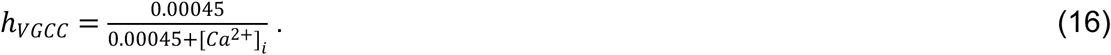

The value of *m*_*VGCC*_ exponentially relaxed to the steady-state value of *m*_*VGCC∞*_:

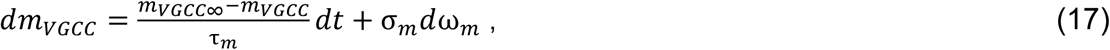

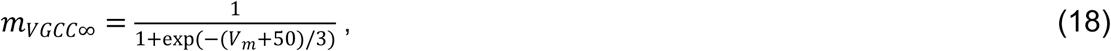

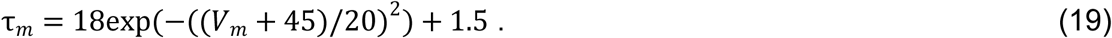

To account for stochastic openings of VGCCs, we added a Wiener process *σ*_*m*_*dω*_*m*_. The variance of the zero-mean Gaussian white noise *σ*_*m*_^*2*^ was defined as in (Riera, Hatanaka, Uchida, Ozaki, & Kawashima, 2011):

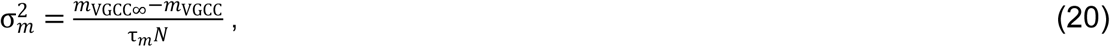

where *N* is the number of VGCCs.

Astrocytic membrane potential (*V*_*m*_) is mostly determined by K^+^ conductance. Therefore, in this study, the K^+^ Nernst potential was used to approximate *V*_*m*_ by adding a modulation factor *ε*, which was chosen from the previous experiments (Schipke et al., 2008):

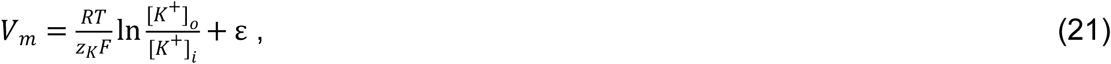

where *R* is the ideal gas constant, *T* is the absolute temperature, *z*_K_ is the valence of K^+^ and *F* is the Faraday constant. [K^+^]_o_ and [K^+^]_i_ are the extracellular and intracellular K^+^ concentrations, respectively.

*E*_*Ca*_ is the Nernst potential for Ca^2+^:

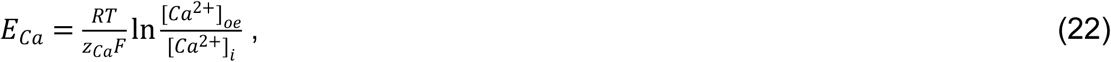

where [*Ca*^2+^]_oe_ is the effective extracellular Ca^2+^ concentration. The compliance with experimental data was achieved in case of [Ca^2+^]_oe_ = (2.00181 + 0.21691 ln([*Ca*^2+^]_o_ - 0.0099)).

The leak efflux from the cytosol to the extracellular space was calculated following (Ullah et al., 2006):

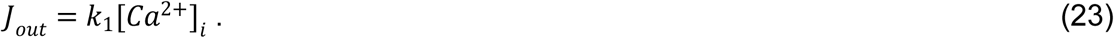

[IP_3_] was determined by the IP_3_ production by phospholipase C (PLC)δ (*J*_*δ*_) and by the IP_3_ degradation as described in (Ullah et al., 2006):

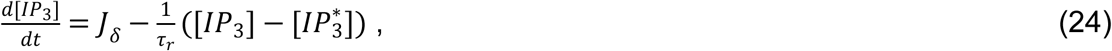

where [IP_3_*] is steady state concentration of IP_3_ and *J*_*δ*_ was modeled as described in (De Young & Keizer, 1992; Ullah et al., 2006):

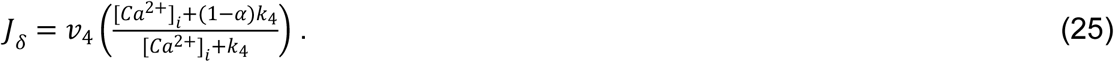

A sketch of the fluxes is shown in Figure 4A. The descriptions and the values of used parameters are given in the Table 1.

## 3. Results

### 3.1 Spontaneous Ca^2+^ events initiation points depend on the width of the process

Ca^2+^ activity was monitored in individual astrocytes transfected with the genetically encoded Ca^2+^ sensor GCaMP2. To avoid interference from neighboring astrocytes, imaging was performed in low-density astrocytic cultures. The morphological profile of individual astrocytes was determined from the baseline fluorescence of GCaMP2 (Figure 1A). The size of the process was defined by the width of the cross-section at half of the maximum fluorescence (half-width; HW) (Figure 1B). 18 cross-sections with 10° increment were plotted at every pixel within the astrocytic territory (Figure 1C). The minimum HW among the 18 cross-sections was then assigned to the pixel (Figure 1D). The astrocyte territory was colored according to the local HWs (Figure 1E,G). The probability density of local HWs (Pr(HW)) had the mode of 1.8 ± 0.3 µm and the median of 4.5 ± 0.3 µm (n = 22 cells) (Figure 1F,H). The result suggests that more than 50% of the territory of a single astrocyte consists of processes with small HW.

Spontaneous Ca^2+^ activity in astrocytes was recorded at 34°C with time-lapse imaging at a speed of 2 frames per second in the presence of Na^+^ channel, mGluR, NMDA, AMPA, GABA_B_, and P2 receptors’ blockers. In agreement with previous reports, conventional region of interest (ROI)-based analysis demonstrated compartmentalization of Ca^2+^ transients (Figure 2A,B) (Di Castro et al., 2011; Haustein et al., 2014; Inagaki, Fukui, Ito, Yamatodani, & Wada, 1991; Shigetomi, Kracun, Sofroniew, & Khakh, 2010). However, this analysis provides very limited spatial information and cannot accurately locate the starting points of Ca^2+^ events. Ca^2+^ transients detected in a particular ROI may not start there, but propagate from other, sometimes, very distant parts of the cell (Agarwal et al., 2017; Heller & Rusakov, 2015; Wu et al., 2014). Therefore, we performed pixel-by-pixel analysis and identified the entire territories of individual Ca^2+^ events in each frame (see Materials and Methods). The volume formed by spatiotemporally inter-connected pixels belonging to the same Ca^2+^ transient was defined as a ‘whole Ca^2+^ event’. Indeed, Ca^2+^ transients detected with ROI-based method represented different parts of whole Ca^2+^ events reconstructed in x - y - time 3D space (Figure 2C,D and Supporting Information Movie 1). To identify the initiation point of each whole Ca^2+^ event, we analyzed the frame in which the event was first detected and calculated the centroid of the detected event area (weighted by ΔF/F amplitude of each pixel, Supporting Information Movie 1). Then each initiation point was assigned to the local HW (Figure 2E). Most Ca^2+^ events started in thin processes (the mode of the event initiation points was at the local HW of 2.5 ± 0.1 µm, n = 22 cells; Figure 2E,F). Notably, the mode of the initiation points coincided with the mode of the local HWs. This opens a possibility that the spontaneous Ca^2+^ events occurred by chance in every part of the astrocyte. If this is the case, the ratio between the probability density of Ca^2+^ event initiation points at the particular HWs and the probability density of the local HWs (Pr(Ca^2+^ events)/Pr(HW)) should be equal to 1. However, Ca^2+^ events occurred in the processes with the HW > 1 µm and < 4 µm with higher probability than the chance level, and in the processes with the HW < 1 µm and > 4 µm with lower probability than the chance level (n = 22; F(1,546) = 545.8, p < 0.0001; interaction: F(25,546) = 17.18, p< 0.0001; two-way RM ANOVA; Figure 2G,H).

### 3.2 SVR of the process determines the probability of Ca^2+^ event initiation

The local HW may not accurately describe the SVR of the process in cultured astrocytes. For example, a very flat process may have a large HW and high SVR. To calculate the actual SVR, we assumed that GCaMP2 is evenly distributed in the cytoplasm and the amount of baseline fluorescence detected with CCD reflects the height of the astrocyte in each pixel. After recording the fluorescence with CCD, the same samples were transferred to the confocal microscope and z-stack was obtained. The actual height calculated from z-stack was used to determine the conversion factor of CCD-recorded fluorescence to process height (Figure 3A-D). A heatmap of SVRs was created for each recorded astrocyte (Figure 3E). The distribution of the local SVRs, had the mode of 0.74 ± 0.04 µm^−1^ and the median of 0.72 ± 0.04 µm^−1^ (n = 22 cells, Figure 3F).

**Figure 3.**
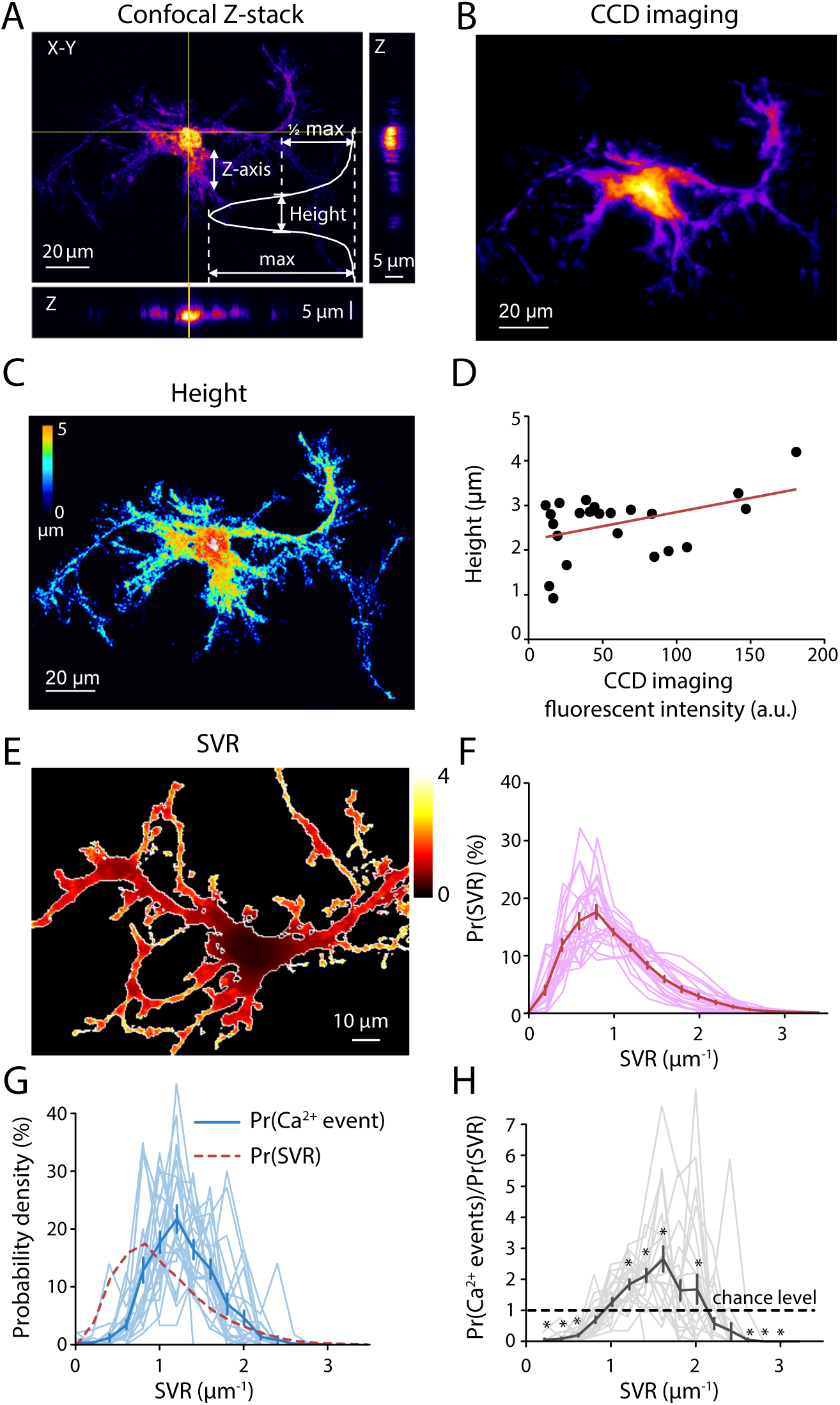
The optimal range of SVRs for Ca^2+^ events initiation in astrocytic processes. **(A)** A maximum projection of a z-stack of the confocal images of a cultured astrocyte expressing GCaMP2. The thickness of the astrocytic processes was approximated by the HW of fluorescence intensity profile (inset). **(B)** The CCD image of baseline fluorescence of GCaMP2 of the astrocyte shown in (A). **(C)** A representative example of the height of the astrocyte in (A) and (B) presented with a heat map. **(D)** The correlation between the height (estimated by confocal imaging) and the fluorescent intensity (acquired by CCD imaging) of the same processes (n = 23 processes/3 cells). **(E)** The local SVR of the astrocyte presented as the color map. Hotter color indicates processes with higher SVR. **(F)** The probability density distribution of SVRs (Pr(SVR)) of astrocytes (thin lines - individual cells, n = 22 cells, thick line mean ± SEM). **(G)** The probability density of Ca^2+^ events initiation points at particular SVRs (Pr(Ca^2+^ events), blue line, thin lines - individual cells, n = 22 cells, thick line mean ± SEM). Dashed pink line indicates Pr(SVR) as shown in (F). **(H)** The summary of the Pr(Ca^2+^ event)/Pr(SVR) ratio (thin lines—individual cells, n = 22 cells, thick line mean ± SEM). The ratio was significantly higher than the chance level in the processes with SVRs ≥ 1.2 µm^−1^ and < 2 µm^−1^. *p < 0.05 ANOVA *post-hoc* Bonferroni’s multiple comparison.

Most Ca^2+^ events started in processes with the SVR > 1 µm^−1^ (the mode of event initiation points was at the local SVR of 1.25 ± 0.07 µm^−1^, n = 22 cells; Figure 3G). Similarly, we calculated the ratio between probability density of Ca^2+^ event starting at particular SVRs and the probability density of SVRs (Pr(Ca^2+^ events)/Pr(SVR)). Ca^2+^ events in the processes with the SVR ≥ 1.2 µm^−1^ and < 2 µm^−1^ occurred with higher probability than the chance level, and in the processes with the SVR ≤ 0.6 µm^−1^ and > 2.4 µm^−1^ lower probability than the chance level (n = 22, F (14, 294) = 17.21, p < 0.0001; interaction: F (14, 294) = 17.21, p < 0.0001; two-way RM ANOVA; Figure 3H).

Our finding suggests that there is a range of the local SVRs in an astrocyte where spontaneous Ca^2+^ events predominantly initiate. Ca^2+^ events have less chance to start in very thin or in thick processes. Absence of Ca^2+^ stores in thin processes can potentially explain the lack of Ca^2+^ event initiation points there (Patrushev, Gavrilov, Turlapov, & Semyanov, 2013). However, this does not rule out the existence of ‘subthreshold’ Ca^2+^ transients, which could not be detected with frame rate or sensitivity of our imaging, in these processes. In the thicker processes, reduced probability of Ca^2+^ events can be explained by the low SVR of these processes. Low SVR suggests fewer Ca^2+^ conductances per volume of the cytosol assuming even surface density of these conductances in the astrocyte. Thus, Ca^2+^ entry through plasma membrane cannot reach the threshold to trigger Ca^2+^ release from endogenous Ca^2+^ stores through IP_3_Rs. This phenomenon is illustrated with a cylindrical model of astrocyte representing a chunk of the astrocytic process (See Materials and Methods, Figure 4A). In this model Ca^2+^ ions stochastically entered the process through the plasma membrane (*J*_*in*_, Figure 4B), leaked from the ER (*J*_*leak*_) and were released from the ER through IP_3_Rs (*J*_*IP3*_). Both Ca^2+^ and IP_3_ were required for activation of the IP_3_Rs as described in (Y. X. Li & Rinzel, 1994). In the presence of IP_3_, the IP_3_Rs were activated by low [Ca^2+^]_i_ and inactivated by high [Ca^2+^]_i_. Ca^2+^ was pumped from the cytosol to the ER by SERCA (*J*_*pump*_) or was moved out of the cell by pumps/exchangers in the plasma membrane (*J*_*out*_). The initial IP_3_ concentration changed due to Ca^2+^-dependent activation of PLCδ (*J*_*IP3*_). *J*_*VGCC*_ depended on a single channel Ca^2+^ conductance, number of channels and Ca^2+^ driving force. Because the density of Ca^2+^ channels was set uniformly on the plasma membrane, the processes with a thicker diameter (lower SVR) had fewer channels per volume of the cytosol. This resulted in smaller fluctuations of baseline [Ca^2+^]_i_ which reached the threshold for IP_3_R activation less frequently. Thus, the frequency of the IP_3_R-mediated Ca^2+^ events was higher in the thinner than in the thicker processes (Figure 4C). At some point the process diameter became smaller than the diameter of the ER (∼1 µm), and such processes could not accommodate the organelle and thus did not maintain IP_3_R-mediated Ca^2+^ events (Figure 4C). Our model reproduced the experimental results showing that the IP_3_R-mediated Ca^2+^ events appear more frequently in a certain range of sizes of astrocytic processes. The model suggests that the local SVR and the presence of the ER in the process are sufficient to explain higher frequency of Ca^2+^ events in distal astrocytic processes without need for any gradients of Ca^2+^ channels in the plasma membrane.

**Figure 4.**
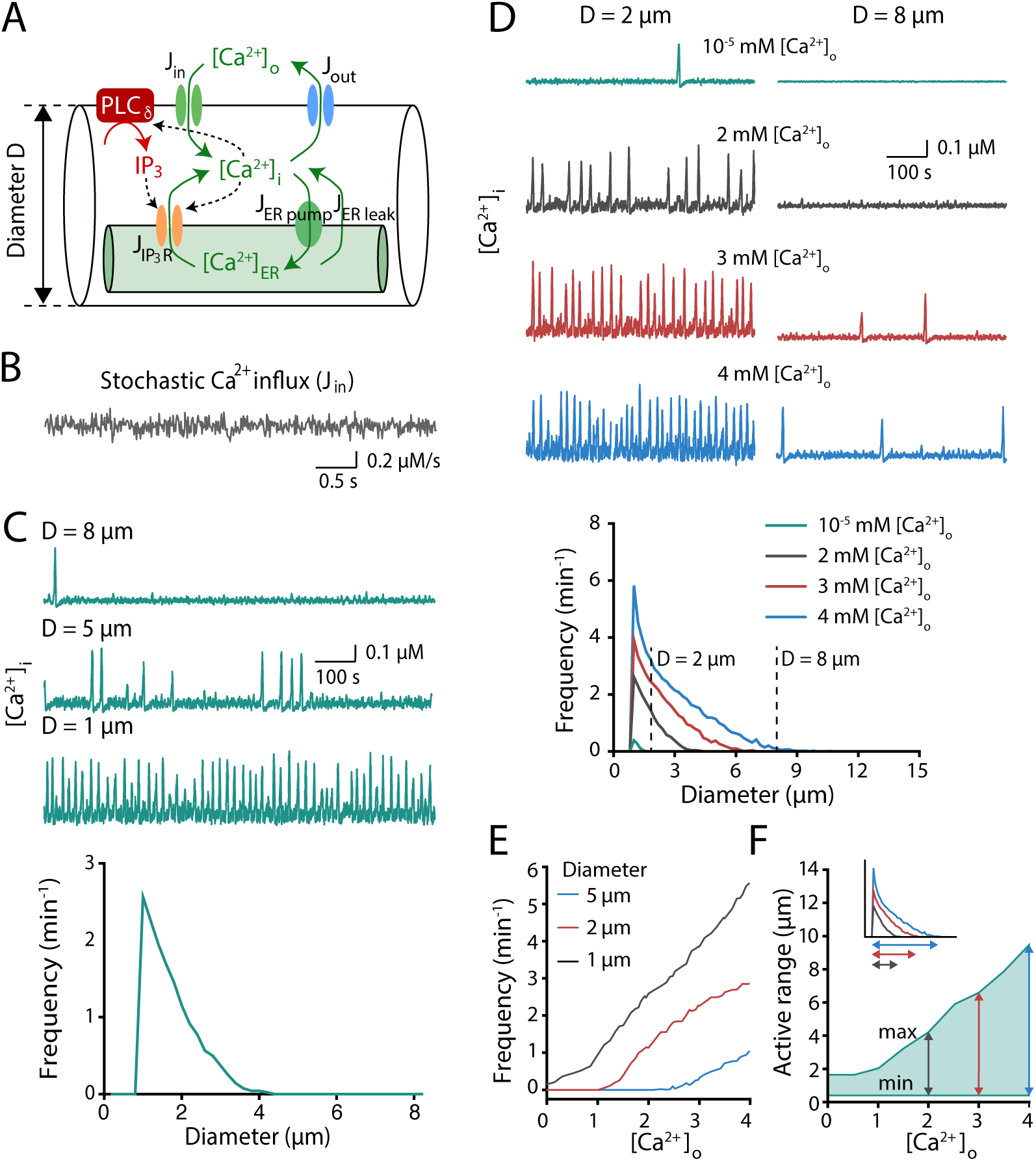
The frequency of spontaneous Ca^2+^ events depends on the diameter of the model astrocytic process and the [Ca^2+^]_o_. **(A)** A schematic of a single compartment model of an astrocytic process. **(B)** Stochastic behavior of modeled Ca^2+^ influx (*J*_*in*_). **(C)** Reducing the diameter of the modeled process increased the frequency of spontaneous Ca^2+^ events. *Top*, representative traces of Ca^2+^ dynamics in the astrocytic processes of three different diameters - *D*. *Bottom*, the frequency of spontaneous Ca^2+^ events as a function of the diameter of the modeled astrocytic process. **(D)** The frequency of the spontaneous Ca^2+^ events is modulated by [Ca^2+^]_o_. *Top*, representative traces of Ca^2+^ dynamics in the astrocytic processes of two different diameters at different [Ca^2+^]_o_: 0.01 µM, 2 mM, 3 mM, and 4 mM. *Bottom*, the frequency of the spontaneous Ca^2+^ events as a function of the diameter of the astrocytic process. **(E)** The dependency of the Ca^2+^ event frequency on [Ca^2+^]_o_ in the astrocytic processes of three different diameters. **(F)** The effect [Ca^2+^]_o_ on the range of astrocytic process diameters where Ca^2+^ events occurred (‘active range’; green-shaded area). *Inset* illustrates how the ‘active range’ was determined in (D). ‘min’: minimum active diameter; ‘max’: maximum active diameter.

### 3.3 Ca^2+^ entry through VGCCs is involved in spontaneous Ca^2+^ activity in cultured astrocytes

In our model, *J*_*VGCC*_ depended on the Ca^2+^ driving force which changes at different concentrations of extracellular Ca^2+^([Ca^2+^]_o_). Indeed, nominally Ca^2+^-free solution ([Ca^2+^]_o_ = 10 µM) suppressed Ca^2+^ events in the processes of all diameters (Figure 4D), while [Ca^2+^]_o_ elevation from 2 mM to 4 mM increased both the peak frequency of Ca^2+^ events and extended the “active range” of HWs where Ca^2+^ events could be triggered (Figure 4E,F). In agreement with the model prediction, removal of extracellular Ca^2+^ suppressed Ca^2+^ events in cultured astrocytes (the effect of HW: F(19,200) = 9.457, p < 0.0001; the effect of [Ca^2+^]_o_ removal: F(1,200) = 26.26, p < 0.0001; interaction: F(19,200) = 5.783, p < 0.0001; n = 11; two-way RM ANOVA; Figure 5) and reduced the active range of HWs (2 mM [Ca^2+^]_o_: 8.7 ± 1.2 µm, 0 mM [Ca^2+^]_o_: 4.8 ± 0.6 µm, n = 11, p < 0.01, Wilcoxon signed-rank test). However, this phenomenon may also be explained by a depletion of Ca^2+^ stores in astrocytes as a result of Ca^2+^ extrusion from astrocytes. Therefore, we gradually increased [Ca^2+^]_o_ from 2 to 4 mM and showed that both the frequency of Ca^2+^ events (the effect of HW: F(19,220) = 5.662, p < 0.0001; the effect of [Ca^2+^]_o_: F(2,240) = 35.48, p < 0.0001; interaction: F(38,240) = 4.825, p< 0.0001; n = 7, two-way RM ANOVA) and the active range of HWs (2 mM [Ca^2+^]_o_: 7.1 ± 1.0 µm, 3 mM [Ca^2+^]_o_: 8.1 ± 1.4 µm, 4 mM [Ca^2+^]_o_: 12.1 ± 1.6 µm, n = 7, p < 0.01, n = 7, Friedman test; Figure 6) were significantly increased. This finding suggests that the transmembrane gradient of Ca^2+^ concentration directly modulates the frequency of spontaneous Ca^2+^ events in astrocytes.

**Figure 5.**
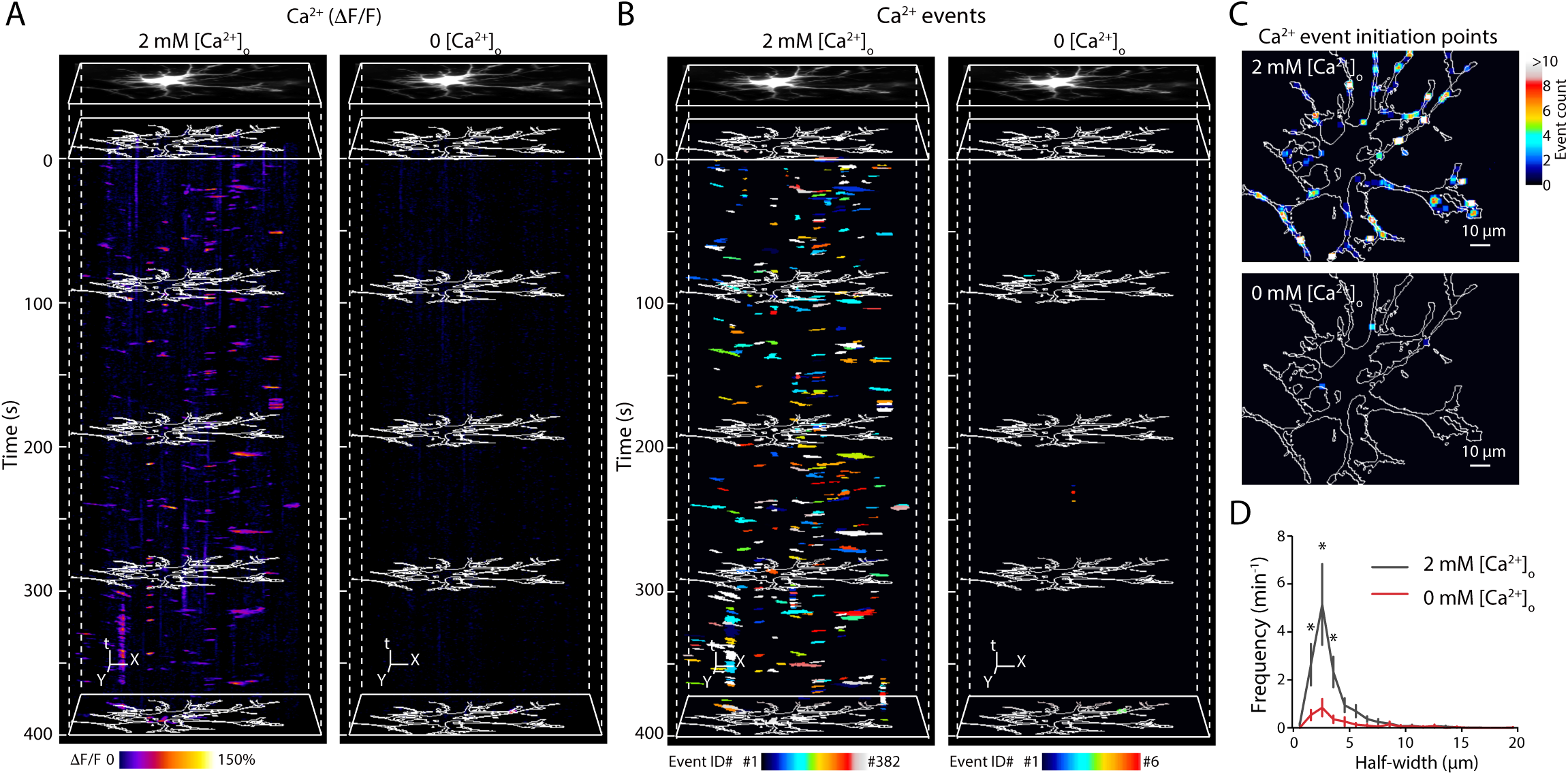
The frequency of astrocytic Ca^2+^ events decreases upon removal of the [Ca^2+^]_o_. **(A)** Representative examples of X-Y-time heat maps of Ca^2+^ dynamics (δF/F) in 2 mM [Ca^2+^]_o_ (*left*) and in 0 mM [Ca^2+^]_o_ (*right*). The white contour shows the boundaries of the astrocyte. **(B)** Same examples as in (A) but presented as X-Y-time binary maps of detected whole Ca^2+^ events. Random colors were assigned to individual events for distinguishing their boundaries. 382 and 6 events were detected during a 400-second recording in 2 mM [Ca^2+^]_o_ and in 0 mM [Ca^2+^]_o_, respectively. **(C)** Representative images of initiation points of Ca^2+^ events in 2 mM [Ca^2+^]_o_ (*top*) and 0 mM [Ca^2+^]_o_ (*bottom*). The centroids of the initial frame of the 382 and 6 detected Ca^2+^ events in 2 mM [Ca^2+^]_o_ and 0 mM [Ca^2+^]_o_, respectively, were plotted and the counts were presented as a color map. **(D)** The summary result of the frequency of event initiation at a given HW (n = 11 cells) in 2 mM [Ca^2+^]_o_ (*black line*) and 0 mM [Ca^2+^]_o_ *(red line*). *p < 0.05 ANOVA *post-hoc* Bonferroni’s multiple comparisons.

**Figure 6.**
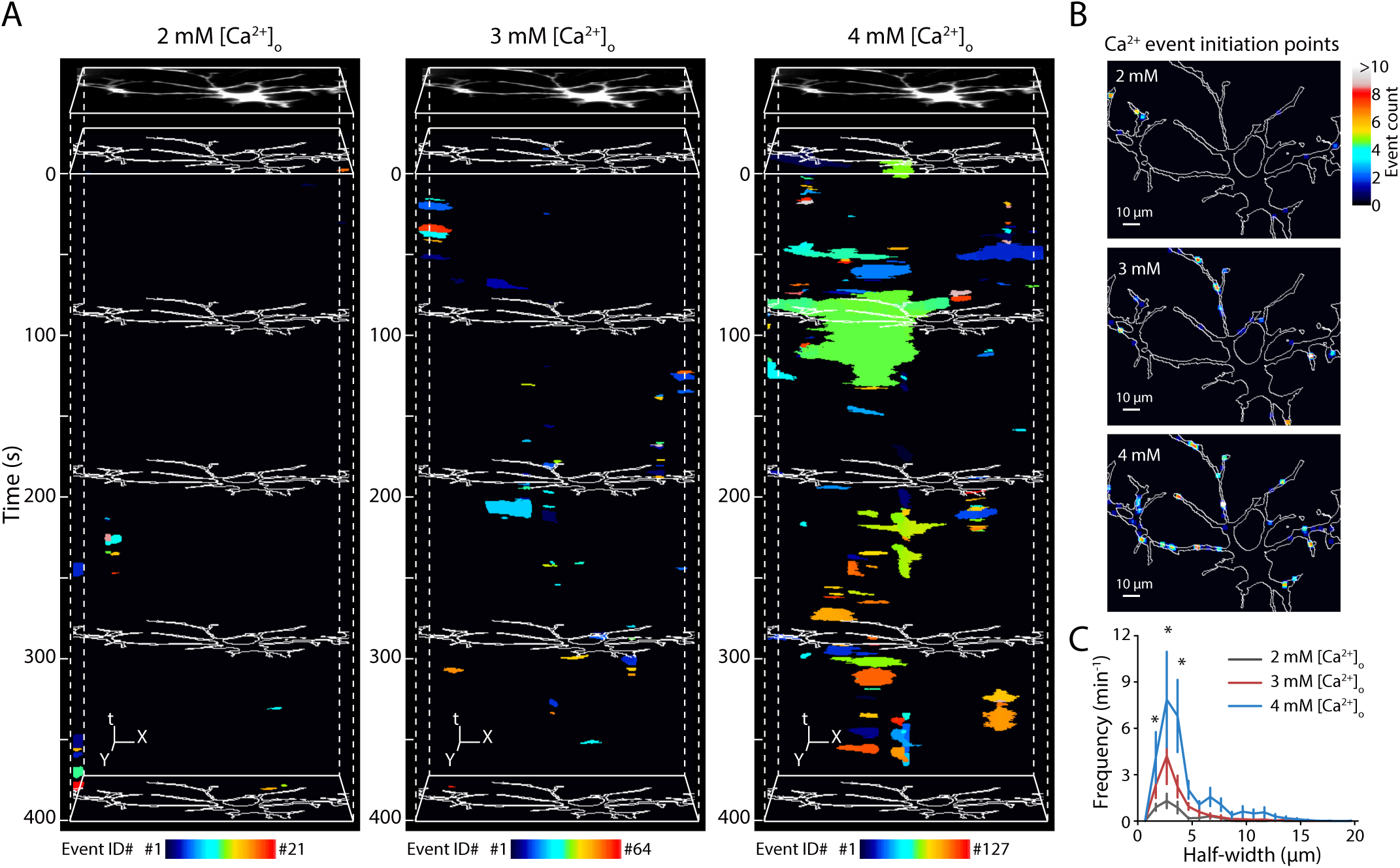
The frequency of astrocytic Ca^2+^ events increases with the [Ca^2+^]_o_ elevation. **(A)** Representative examples of X-Y-time binary maps of detected whole Ca^2+^ events in 2 mM, 3 mM, and 4 mM [Ca^2+^]_o_. Random colors were assigned to individual events. 21, 64, and 127 events were detected in 2 mM, 3 mM, and 4 mM [Ca^2+^]_o_, respectively, during each 400-second recording. The white contour shows the boundaries of the astrocyte. **(B)** Representative images of initiation points of Ca^2+^ events in 2 mM, 3 mM, and 4 mM [Ca^2+^]_o_. The centroids of the initial frame of the 21, 64, and 127 detected Ca^2+^ events in 2 mM, 3 mM, and 4 mM [Ca^2+^]_o_, respectively, were plotted and the counts were presented as color maps. **(C)** The summary result of the frequency of event initiation at a given HW (n = 7 cells) in 2 mM (*black line*), 3 mM (*red line*) and 4 mM (*blue line*) [Ca^2+^]_o_. *p < 0.05 ANOVA *post-hoc* Bonferroni’s multiple comparisons.

Besides [Ca^2+^]_o_, another parameter that determines the transmembrane Ca^2+^ flow is the Ca^2+^ membrane conductance of astrocytic plasma membrane. When neuronal network activity is high, activity-dependent changes in extracellular K^+^ concentration can depolarize the astrocyte and increase the open probability of VGCCs (Bellot-Saez, Kekesi, Morley, & Buskila, 2017; Duffy & MacVicar, 1994). Therefore, we simulated how changes in membrane potential (V_m_) can affect the frequency of Ca^2+^ events in the processes of different diameter. The membrane depolarisation from −80 mV to −64 mV increased both the peak frequency of Ca^2+^ events and extended the range of diameters where Ca^2+^ events were triggered (Figure 7 A,B). To test this prediction experimentally, we increased [K^+^]_o_ from 2 mM to 5 mM. Such changes of [K^+^]_o_ occur in physiological range (Bellot-Saez et al., 2017). Indeed, we observed an increase in the frequency of Ca^2+^ events in thin astrocytic processes (the effect of HW: F(19,160) = 20.67, p < 0.0001; the effect of [K^+^]_o_: F(1,160) = 8.863, p = 0.0034; interaction: F(19,160) = 2.568, p = 0.0007; n = 9, two-way RM ANOVA; Figure 7 C-E) but no significant increase in the active range of HWs (p = 0.88, n = 9, Wilcoxon signed-rank test). When we repeated the same experiment in the presence of VGCC blockers (10 µM mibefradil and 20 µM nimodipine), the increase of Ca^2+^ events frequency was no longer observed (the effect of HW: F(19,100) = 13.53, p < 0.0001; the effect of [K^+^]_o_: F(1,100) = 0.1347, p = 0.7144; interaction: F(19,100) = 0.651, p = 0.8571; n = 6, two-way RM ANOVA; Figure 7F). This finding suggests that the frequency of spontaneous Ca^2+^ event in thin astrocytic processes can be increased by extracellular K^+^ accumulation during local neuronal activity, through activating of VGCCs in the plasma membrane of astrocytes.

**Figure 7.**
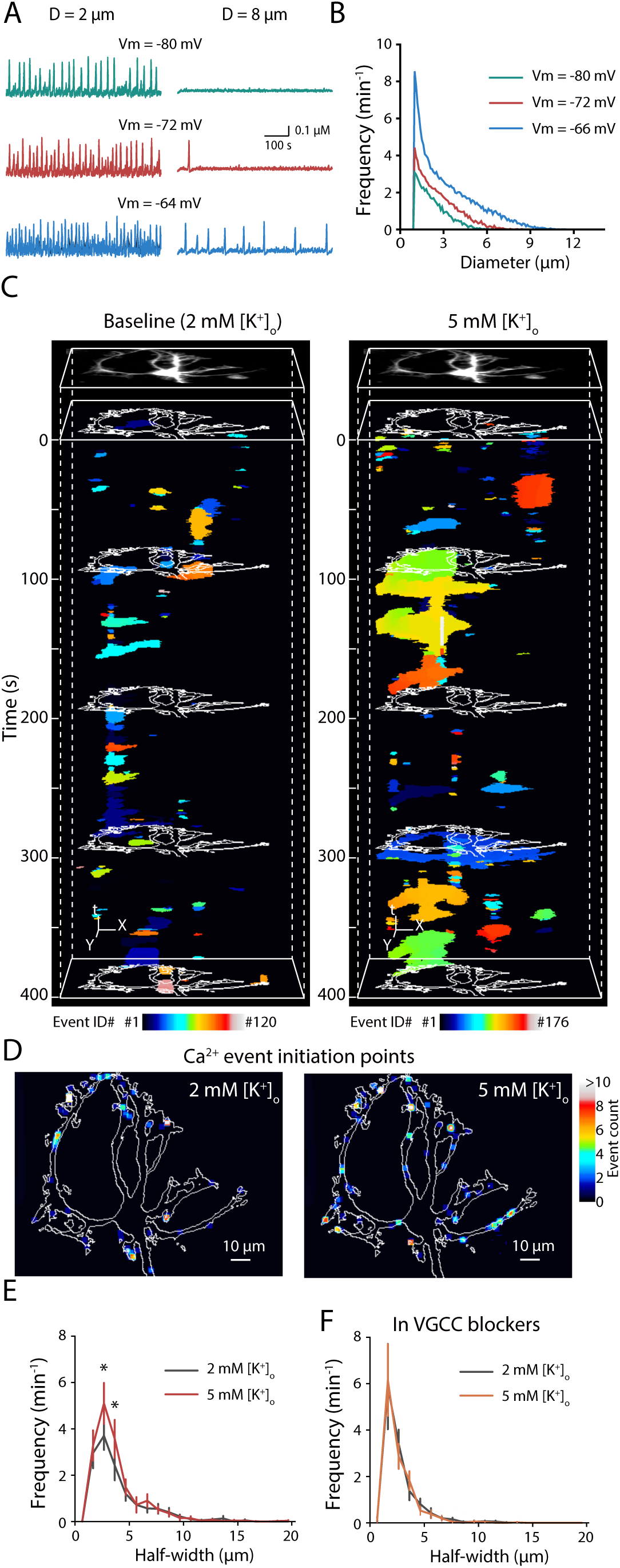
VGCC-dependent increase in the frequency of Ca^2+^ events caused by elevated [K^+^]_o_. **(A-B)** The frequency of spontaneous Ca^2+^ events can be modulated by resting membrane potentials in the modeled astrocyte. **(A)** Representative traces of Ca^2+^ dynamics at three different resting membrane potentials (−80, −72, and −64 mv). **(B)** The frequency of spontaneous Ca^2+^ events plotted as a function of the diameter of the modeled astrocytic process. **(C)** Representative examples of X-Y-time binary maps of detected Ca^2+^ events in the baseline (2 mM [K^+^]_o_), and at 5 mM [K^+^]_o_. Random colors were assigned to individual events for distinguishing their boundaries. 120 and 176 events were detected in 2 mM and 5 mM [K^+^]_o_, respectively, during each 400-second recordings. The white contour shows the boundaries of the astrocyte. **(D)** Representative images of initiation points of Ca^2+^ events in 2 mM and 5 mM [K^+^]_o_. The centroids of the initial frame of the 120 and 176 detected Ca^2+^ events in 2 mM and 5 mM [K^+^]_o_, respectively, were plotted and the counts were presented as color maps. **(E)** The summary result of the frequency of event initiation at a given HW (n = 9 cells) in 2 mM (*black line*) and 5 mM (*red line*) [K^+^]_o_. *p < 0.05 ANOVA *post-hoc* Bonferroni’s multiple comparisons. **(F)** The summary result of the frequency of event initiation at a given HW (n = 6 cells) in 2 mM (*black line*) and 5 mM (*red line*) [K^+^]_o_ in the presence of VGCC blockers.

### 3.4 The role of TRPA1 channels and mitochondria in spontaneous Ca^2+^ activity

TRPA1 channels are also suggested to regulate astrocyte resting Ca^2+^ levels (Shigetomi et al., 2013; Shigetomi, Tong, Kwan, Corey, & Khakh, 2012). However, we did not observe a significant effect of TRPA1 blocker (20 µM HC 030031) on the frequency Ca^2+^ events (the effect of HW: F(19,57) = 5.577, p < 0.0001; the effect of HC 030031: F(1,3) = 0.0126, p = 0.92; interaction: F(19,57) = 0.3351, p = 0.9947; n = 4, two-way RM ANOVA; Supporting Information Figure 1). This finding does not rule out a possibility of Ca^2+^ entry through TRPA1 channels which plays a role in the nearest submembrane space.

A recent report suggests that mitochondrial permeability transition pore may transiently open and induces microdomain Ca^2+^ Transients in astrocytic processes (Agarwal et al., 2017). If this is the case, Ca^2+^ events initiation points should co-localize with mitochondria. To test this hypothesis, we stained the mitochondria with 50 nM MitoTracker Red CMXRos red fluorescent dye and studied their distribution within individual astrocytes (Supporting Information Figure 2A and 2B). If the higher Ca^2+^ event initiation frequency is contributed mainly by mitochondrial Ca^2+^ release, we should expect a higher probability of mitochondrial presence (Pr(Mitochondria)) in processes with small HW (Supporting Information Figure 2C). However, Pr(Mitochondria) was significant lower than Pr(HW) in small processes and larger than Pr(HW) in thicker processes (the effect of HW: F(19,114) = 16.65, p < 0.0001; Pr(Mitochondria) vs Pr(HW): F(1,6) = 1.384, p = 0.28; interaction: F(19,114) = 8.422, p < 0.0001; n = 7, two-way RM ANOVA, *post-hoc* Bonferroni’s multiple comparison; Supporting Information Figure 2C). This finding suggested that even though mitochondria may contribute to astrocytic Ca^2+^ activity (Agarwal et al., 2017), it does not explain the morphology-dependent Ca^2+^ activity we observed here.

### 3.5 Ca^2+^ events start in thin distal process in astrocytes in hippocampal slices

These experiments were done with cultured astrocytes. This preparation allows to image Ca^2+^ dynamics in an entire astrocyte and to measure the HWs/SVRs of the processes. However, astrocytic culture has major drawbacks, since the cultured astrocytes do not develop complex structure characteristic to astrocytes *in vivo* and may express non-specific receptors and channels (Lange, Bak, Waagepetersen, Schousboe, & Norenberg, 2012). Therefore, we performed imaging in hippocampal slices from transgenic animals with astrocyte specific expression of GCaMP2 (Figure 8A). Because fine astrocytic processes were beyond optical resolution and their HWs/SVRs cannot be precisely measured, we estimated the volume fraction (VF) of fine astrocytic processes as the ratio between their fluorescence and the fluorescence of the soma (Medvedev et al., 2014; Plata et al., 2018). Multiple fluorescence profiles were plotted across the soma in each direction and the entire astrocyte map of VFs was generated (Figure 8B-D). Notably, the distribution of VFs in astrocytes in slices resembled the distribution of HWs/SVRs in cultured astrocytes (the mode: 1.41 ± 0.31 % and the median 6.07 ± 0.66 %; n = 8 cells; Figure 8E).

**Figure 8.**
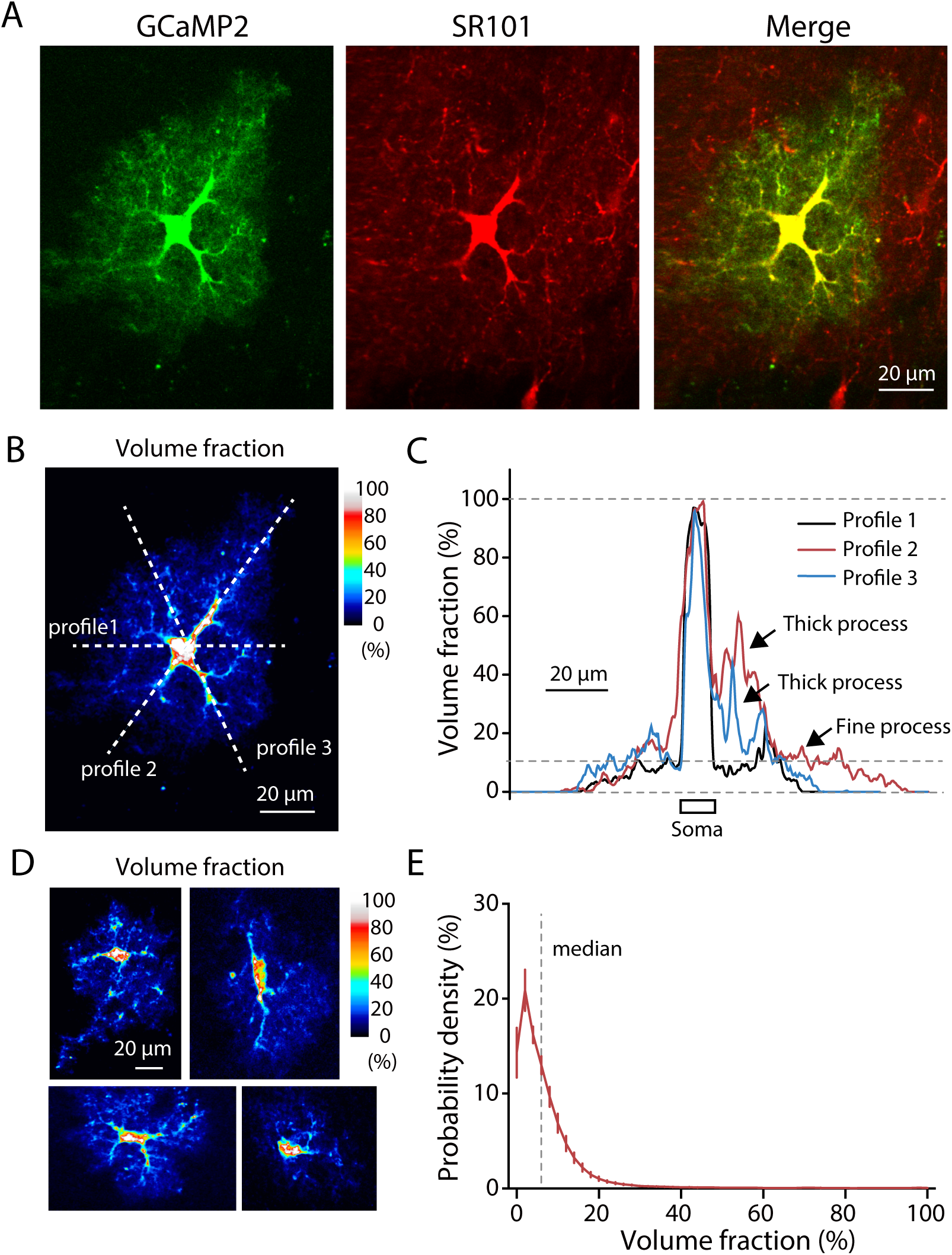
Morphological profile of astrocytes in hippocampal slices. **(A)** A typical astrocyte expressing the Ca^2+^ sensor GCaMP2 in hippocampal slice. *Left,* the baseline fluorescence of GCAMP2; *middle*, astrocyte-specific marker, sulforhodamine 101 (SR101); *right*, merged image of GCaMP2 and SR101. **(B)** A representative heat map of VF of the astrocyte shown in (A). **(C)** Three examples of line profiles (3 dashed lines in (B)) of VF aligned to the soma. The VF was used to characterize unresolved astrocytic processes. **(D)** Representative examples of VF heat map. **(E)** The mean Pr(VF) of several astrocytes (n = 8 cells). The dashed line indicates the median (VF: 6.07 ± 0.66 %) of VF distribution.

We identified the initiation point of each whole Ca^2+^ event (Figure 9A,B; Supporting Information Movie 2). Each initiation point was linked to the local VF (Figure 9C). In agreement with the results obtained in cultured astrocytes, most of Ca^2+^ events started in the sites with low local VF (the mode of event initiation points was at the sites with the VF of 9.06 ± 1.75 %, n = 8 cells; Figure 9C). In contrast to the cultured astrocytes, the mode of the initiation points did not precisely coincide with the mode of the local VF. This may be because the mode of the local VF corresponded to the finest astrocytic processes, i.e. ‘leaflets’. Since the leaflets are devoid of endogenous Ca^2+^ stores they may not initiate the spreading Ca^2+^ events analyzed here (Gavrilov et al., 2018; Patrushev et al., 2013).

**Figure 9.**
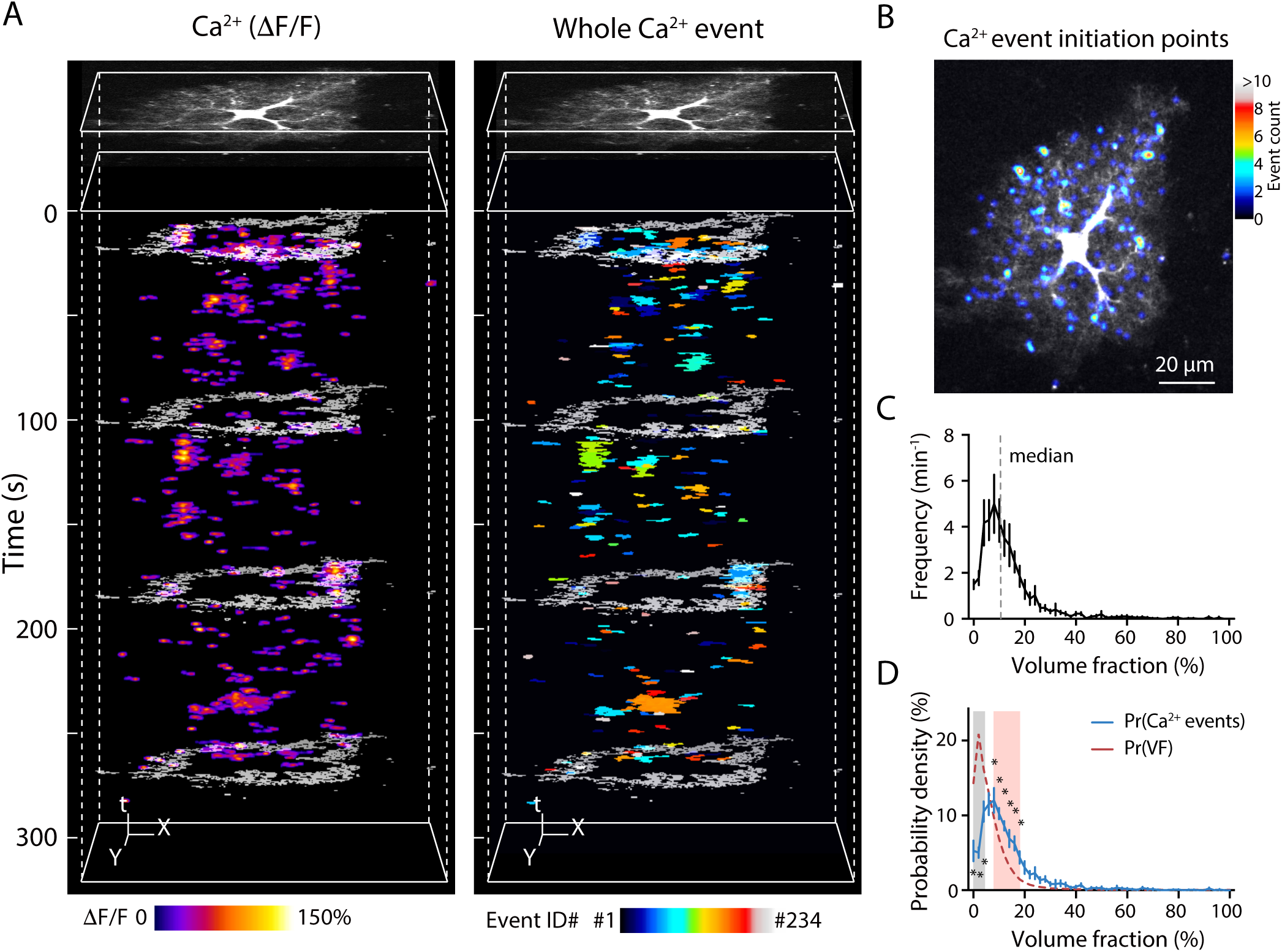
Ca^2+^ events predominantly start in the parts of astrocytic domain with low astrocytic VF. **(A)** *Left*, a representative example of X-Y-time heat map of Ca^2+^ dynamics (δF/F) of detected events. The white contour shows the boundaries of the astrocyte. *Right*, a representative example of X-Y-time binary map of detected whole Ca^2+^ events. Random colors were assigned to individual events to distinguish their boundaries. **(B)** A representative image of event counts of Ca^2+^ event initiation points (*color map*) overlapping with the baseline GCaMP2 fluorescence (*grey scale*). **(C)** The mean Ca^2+^ events frequency plotted against the VF of the events initiation points (n = 8 cells). **(D)** Comparison of Pr(Ca^2+^ events) (*blue line*) with Pr(VF) (*red dash line* re-plotted from Figure 8E) (n = 8 cells). The Pr(Ca^2+^ events) was significantly higher than Pr(FV) in thinner processes of VF = 8 - 18 % (*red shaded area*) but significantly lower in the finest processes of VF = 0 – 6 % (*grey shaded area*). *p < 0.05 ANOVA post-hoc Bonferroni’s multiple comparisons.

## 4. Discussion

Our findings demonstrate that the initiation points of spontaneous Ca^2+^ events are predominately located in thin processes within an optimal range of local HWs/SVRs in cultured astrocytes, and within an optimal range of local VFs within astrocytic domain in hippocampal slices. By combining astrocytic Ca^2+^ imaging with computational simulation, we show that higher SVR of thin astrocytic processes determines higher frequency of spontaneous Ca^2+^ events. Larger fluctuation of [Ca^2+^]_i_ occur within limited volume of the cytosol due to stochastic Ca^2+^ entry through the plasma membrane. Some of such fluctuations reach the threshold for the activation of IP_3_Rs, which require Ca^2+^ as a co-agonist along with IP_3_ (Bezprozvanny, Watras, & Ehrlich, 1991; Dupont & Goldbeter, 1993; Shinohara et al., 2011). Subsequent Ca^2+^ release from the ER amplifies the suprathreshold Ca^2+^ fluctuations, triggering longer and larger Ca^2+^ events detectable with full-frame time-lapse imaging. However, such Ca^2+^ events do not start in astrocytic leaflets that are devoid of Ca^2+^ stores (Patrushev et al., 2013). This finding do not rule out that fast Ca^2+^ sparks occur in these compartments near the plasma membrane because of direct Ca^2+^ entry (Rungta et al., 2016; Shigetomi et al., 2010). Such sparks may not be detectable in our experimental conditions (resolution and speed of imaging) but still could play a role in astrocyte physiology, e.g. trigger a local release of gliotransmitters. A recent computational study has also suggested that local Ca^2+^ sparks can be amplified through NCX (Brazhe et al., 2018). Moreover Ca^2+^ sparks in ER/mitochondria-free leaflets can potentially integrate to trigger spreading Ca^2+^ events in their parent astrocytic branches containing the Ca^2+^ stores. This scenario is reminiscent of excitatory postsynaptic potentials (EPSPs) integration and triggering of action potentials in neurons. EPSP amplitude depends both on the number of activated receptors and on the cell input resistance (Blomfield, 1974; Pavlov et al., 2014; Song, Savtchenko, & Semyanov, 2011). Higher input resistance makes EPSPs larger and increases their chance to trigger an action potential. Similarly, higher SVR of the astrocytic process makes the amplitude of [Ca^2+^]_i_ fluctuations larger and increases their chance to trigger CICR from endogenous Ca^2+^ stores. This notion is also supported by the existence of a [Ca^2+^]_i_ gradient in astrocytes: [Ca^2+^]_i_ is larger in thin distal processes than in thick proximal processes or in soma (Zheng et al., 2015). Thus, astrocytic Ca^2+^ activity may be directly regulated by changes in the morphology of astrocytic processes which occur both in physiological and pathological conditions (Heller & Rusakov, 2015). Retraction of astrocytic leaflets and redistribution of glial fibrillary acidic protein (GFAP) has been reported in astrocytes of supraoptic nucleus during the hormonal changes associated with lactation (Salm, Smithson, & Hatton, 1985). A decreased astrocytic presence in the synaptic microenvironment promotes glutamate spillover in lactating animals (Oliet, Piet, & Poulain, 2001). In contrast, synaptic long-term potentiation (LTP) is associated with increased density of astrocytic leaflets around potentiated synapse (Bernardinelli, Muller, & Nikonenko, 2014; Bernardinelli, Randall, et al., 2014; Wenzel, Lammert, Meyer, & Krug, 1991). Chronic whisker stimulation also increases astrocytic coverage of synapses in the barrel cortex (Genoud et al., 2006).

Striking changes in morphology of astrocytic processes has been observed during aging (Rodriguez et al., 2014). Age-related changes are complex and region-specific: astrocytes become hypertrophic in hippocampus, while lose processes in the entorhinal cortex. Morphological changes are accompanied by corresponding changes in astrocytic Ca^2+^ signaling (Lalo, Palygin, North, Verkhratsky, & Pankratov, 2011). Astrocytic atrophy parallels reduced Ca^2+^ activity in hippocampal astrocytes in the pilocarpine model of epilepsy (Plata et al., 2018). Astrocytic atrophy has been observed in a transgenic mouse model of Alzheimer’s disease (3xTg-AD) in hippocampus as well as entorhinal and prefrontal cortices (Rodriguez, Jones, & Verkhratsky, 2009). The effect of astrocytic atrophy in 3xTg-AD mice reverses when the animals are subjected to physical activity or are kept in an enriched environment (Rodriguez, Terzieva, Olabarria, Lanza, & Verkhratsky, 2013). All these lines of evidence suggest that astrocytic processes are highly plastic, and the proportion of branches and leaflets can rapidly change. However, in most cases such changes in astrocytic morphology have been considered in the context of the spatial relationship between astrocytic processes and synapses. Our data suggest that morphological reorganization of astrocytes can be directly translated into changes of their Ca^2+^ activity. Indeed, an increase in complexity of astrocytic processes during juvenile development parallels an increase in spread of subcellular Ca^2+^ events in mouse hippocampal astrocytes (Nakayama et al., 2016).

Although our results suggest that a higher SVR alone can explain a higher probability of Ca^2+^ events in thin processes, they do not rule out potential contribution of subcellular gradients of receptors, channels and transporters. When we varied [Ca^2+^]_o_ we have observed corresponding changes in the Ca^2+^ events frequency. The increase in [Ca^2+^]_o_ produced the proportional increase of Ca^2+^ events in thin and thick processes, suggesting a uniform distribution of Ca^2+^ conductances. In contrast, [K^+^]_o_ elevation produced VGCC-dependent increase in the frequency of Ca^2+^ events only in thin astrocytic processes. This is consistent with a higher density of either K_ir_ (responsible for the cell depolarization) or VGCC in thin astrocytic processes (Verkhratsky & Steinhauser, 2000). Notably, Ca^2+^ dynamics in astrocytes is strongly influenced by temperature (Komin, Moein, Ellisman, & Skupin, 2015; Schipke et al., 2008). Here we performed the experiments at 34°C. However, a previous report suggests that Ca^2+^ events in astrocytes become more frequent and slower at the room temperature (Schipke et al., 2008). This effect is linked to a decreased strength of SERCA (Komin et al., 2015). As a result, SERCA can pump Ca^2+^ to ER less efficiently and therefore Ca^2+^ remains longer in the cytosol. Thus, SERCA strength regulates both the frequency of Ca^2+^ events and their durations.

One of the major drawbacks of experiments performed in cultured astrocytes is that the cells do not develop in a physiological environment. Cultured astrocytes are morphologically very different and can express receptors and channels which are not characteristic to astrocytes *in vivo* (Lange et al., 2012). However, the use of cultured astrocytes in the current study was justified. We could image Ca^2+^ activity within the ‘domain’ of entire astrocyte and could resolve astrocytic processes using light microscopy. Our conclusions are related purely to astrocyte geometry and can further extrapolated to thinner astrocytic processes found *in vivo*. To confirm this, we performed two-photon imaging experiments in hippocampal slices of transgenic mouse with astrocyte-specific expression of GCaMP2. Fine astrocytic processes could not be resolved with diffraction-limited optical imaging. To circumvent this problem, we analyzed the local VF of astrocytic processes (Medvedev et al., 2014; Plata et al., 2018). This measure reflects both the thickness and the density of local astrocytic processes. Nevertheless, this approach gave very similar result to the cultured astrocytes: most of the astrocyte’s domain is occupied by the territory with low VF and most of the Ca^2+^ events are triggered within an optimal range of these VFs. Another potential caveat of Ca^2+^ imaging in slices was that the initiation points of Ca^2+^ events may be located outside of the focal plane. This issue can be addressed in the future experiments with emerging technologies such as three-dimensional Ca^2+^ imaging (Savtchouk, Carriero, & Volterra, 2018).

## Acknowledgments

The authors thank Dr. Hiroko Bannai for help with primary hippocampal astrocyte/neuron co-culture and transfection. The research was supported by the Russian Science Foundation (project No. 16-14-00201).

